# Development of immobilized antibody-based affinity grid strategy and application in on-grid purification of target proteins for high-resolution cryo-EM

**DOI:** 10.1101/2024.01.17.575977

**Authors:** Qiaoyu Zhao, Xiaoyu Hong, Yanxing Wang, Shaoning Zhang, Zhanyu Ding, Xueming Meng, Qianqian Song, Qin Hong, Wanying Jiang, Xiangyi Shi, Tianxun Cai, Yao Cong

**Affiliations:** Key Laboratory of RNA Science and Engineering, Shanghai Institute of Biochemistry and Cell Biology, Center for Excellence in Molecular Cell Science, Chinese Academy of Sciences, 200031, China; University of Chinese Academy of Sciences, Shanghai; State Key Laboratory of High Performance Ceramics and Superfine Microstructure, Shanghai Institute of Ceramics, Chinese Academy of Sciences, Shanghai 200050; Key Laboratory of Systems Health Science of Zhejiang Province, School of Life Science, Hangzhou Institute for Advanced Study, University of Chinese Academy of Sciences, Hangzhou, China; Shanghai Nanoport, Thermofisher Scientific, Shanghai, China

**Keywords:** immobilized antibody-based affinity grid, on-grid purification, preferred orientation, atomic resolution cryo-EM

## Abstract

In cryo-electron microscopy (cryo-EM), sample preparation, especially for rare or fragile macromolecular assemblies and those suffering from air-water interface denaturation and particle orientation distribution problems, is a major bottleneck. Here, we developed and characterized an immobilized antibody-based affinity grid (IAAG) strategy based on the high-affinity PA tag/NZ-1 antibody epitope tag system. We used Pyr-NHS as a linker to immobilize NZ-1 Fab on the graphene oxide or carbon covered grid surface. We showed that the IAAG grid can enrich the PA-tagged target proteins, and overcome preferred orientation problems. Furthermore, we demonstrated that our IAAG strategy can be utilized for on-grid purification of low-abundance target complexes from cell lysates and enables atomic resolution cryo-EM. This approach greatly streamlines the purification process, reduces the need for large quantities of biological samples, and addresses common challenges encountered in cryo-EM sample preparation. Collectively, our IAAG strategy provides an efficient and robust means for combined sample purification and vitrification feasible for high-resolution cryo-EM. This approach also has the potential for broader applicability in cryo-EM and cryo-ET.

## Introduction

With the rapid progresses in instrumentation and data analysis methods/software, cryo-electron microscopy (cryo-EM) single-particle analysis has emerged as a powerful tool for the structural characterization of biomolecules, particularly macromolecular complexes^1–6^. However, for low abundance and/or fragile macromolecular complexes, as well as biomolecules exhibiting preferred orientation, disintegration or aggregation problems related to air-water interface (AWI) interference, purification and vitrification process has become a bottleneck for cryo-EM^7–12^. A promising approach involves the utilization of affinity grids for on-grid specimen purification^13–17^. While initial applications have been limited in scope and resolution, with most resolutions limited to 8-40Å^13,15,16,18–20^ (with a recent exception of MS2 virions at a resolution of 3.03 Å^17^), it is conceivable that a highly specific and well-conjugated tag, in combination with a stringent on-grid purification protocol, could serve as a powerful method to isolate and examine rare biological assemblies^21–23^.

The affinity grid strategy modifies the surface of TEM grids with an affinity layer. This layer extracts and concentrates target biological assemblies on the grid, keeping the target away from the air-water interface. With constant efforts over the past decade, several strategies have been developed for generating affinity layers on TEM grids. These strategies include: (1) Functionalized lipids monolayers. They are functionalized with Ni-NTA or high-affinity small molecule, thereby can capture His-tagged or other target proteins^13,14,20,24,25^. (2) Streptavidin 2D crystals. These can capture biotinylated proteins^26–28^. This strategy has recently been applied in G-quadruplex RNA-mediated PRC2 dimer formation and cryo-EM structure determination^29^. These two strategies require special handling and/or tools as well as user experience for grid preparation, and introduce significant background noise. The later one requires computational removal of the 2D lattice in Fourier space. (3) Antibody-based affinity grids. These concentrate target biomolecules on grid^16,30,31^. However, the antibodies are placed on the grid surface without chemical immobilization, which may not sustain multiple rounds of washing to remove contaminations. (4) Functionalized carbon^32,33^, graphene^34,35^, and graphene oxide (GO) surfaces^15,17,36,37^. These can introduce bioactive ligands onto supporting-film covered cryo-EM grid, enabling the enrichment of target proteins^15,17,32,33,36,37^ or overcome preferred orientation problems^34,35^. However, this strategy requires freshly homemade carbon/graphene/GO supporting films conjugated with sufficient oxygen contents to chemically decorate the surface^15,17,32,34–37^, or customized chemical reagents to introduce biological ligands^15,33,36,37^. Despite these advancements, a more efficient and robust cryo-EM affinity grid strategy is needed to perform on-grid purification, especially for low-abundance or fragile target complexes, and to handle preferred orientation problems.

In a previous study, a novel PA–NZ-1 epitope tag system was developed^38^, which consists of the NZ-1 antibody possessing high affinity toward the dodecapeptide dubbed PA tag, and is characterized by slow dissociation kinetics. This system has been used in subunit identification within macromolecular complexes^39–43^. Leveraging the PA–NZ-1 affinity system, we developed an immobilized antibody-based affinity grid (termed IAAG) strategy for high-resolution cryo-EM. We used 1-Pyrenebutyric acid N-hydroxysuccinimide ester (Pyr-NHS) as a linker to immobilize the antibody or Fab fragment on the grid surface. Our IAAG strategy has been proven effective in concentrating low-abundance complexes and overcoming preferred orientation problems. It is also convenient to apply, without the need for customized GO or chemical reagents. Our study demonstrated that the IAAG strategy can be used in on-grid purification of apoferritin and the CCT6 homo-oligomeric ring (CCT6-HR) complex from cell lysates, facilitating the atomic resolution cryo-EM reconstructions. Overall, the IAAG strategy shows promise for on-grid purification and cryo-EM study of challenging macromolecular complexes at atomic resolution.

## Results

### IAAG strategy development

In our approach, we employ Pyr-NHS as a linker to immobilize NZ-1 Fab to GO/carbon grid surfaces. Pyr-NHS is a chemical crosslinker that can noncovalently anchor onto carbon/graphene/GO covered grid surfaces (sharing sp^2^-hybridized carbon lattice) via π-π interaction through its pyrene moiety. This retains the sp^2^ lattice of the carbon skeleton without potentially disrupting the surface structure and electrical conductivity^44,45^. Concurrently, Pyr-NHS has a bioactive ester head that can react with primary amines in the NZ-1 Fab or other antibody/biomolecules, forming stable amide bonds to immobilize them on the grid surface (Fig. 1A). Pyr-NHS has been used in efficient graphene-based biosensor systems^46–51^. For efficient affinity grid preparation, we opted for commercially available GO coated cryo-EM grid (abbreviated as GO grid) to assemble our IAAG grid. Furthermore, we used only the Fab fragment of the NZ-1 antibody to reduce background noise.

**Fig. 1.**
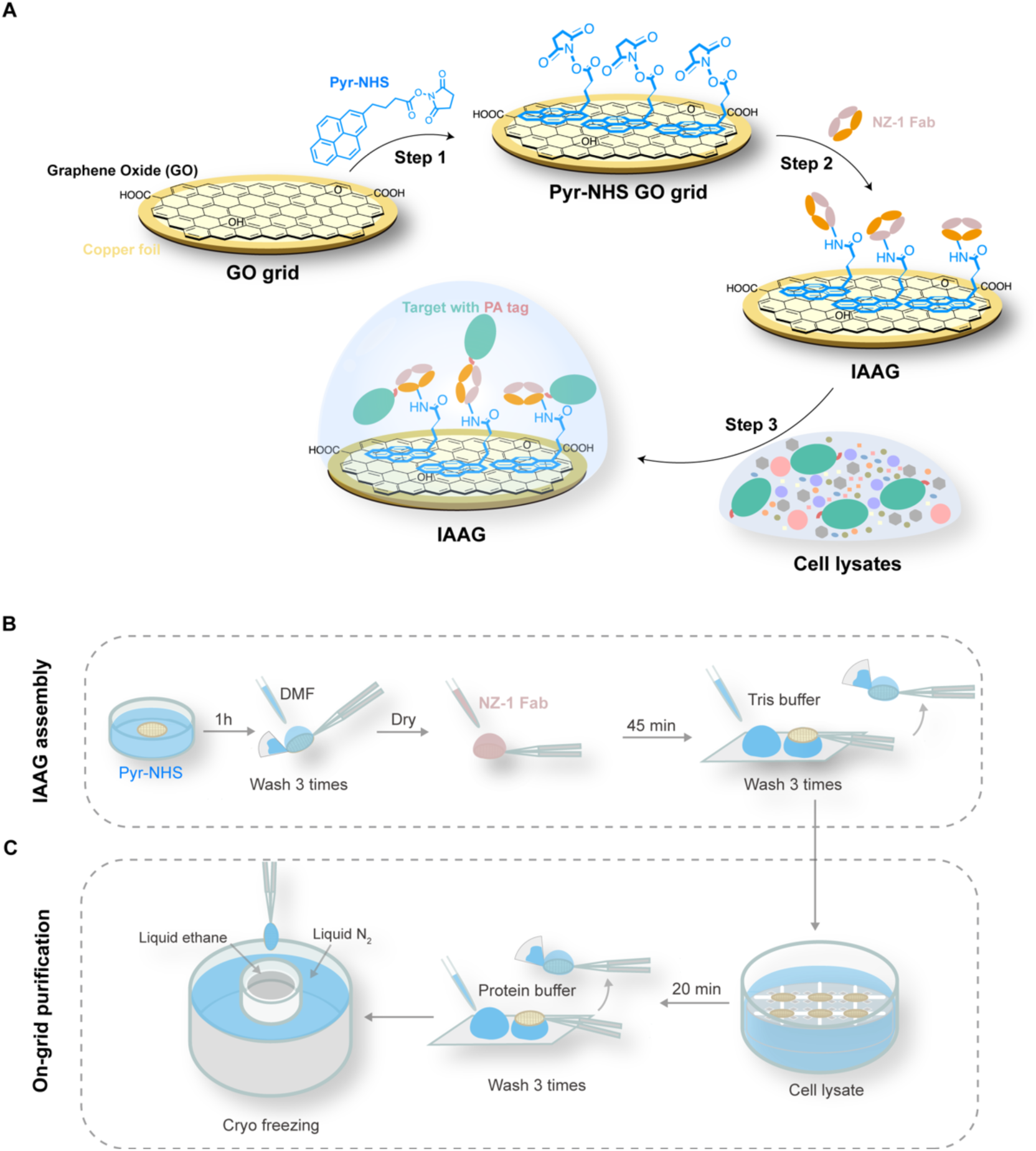
Schematic illustration of the IAAG strategy. (A) Schematic illustration of the assembly of the IAAG grid and the on-grid purification of target proteins for cryo-EM. (B) Workflow for the assembly of the IAAG grid. First, the GO-covered cryo-EM grid is immersed in Pyr-NHS soluted in DMF for 1 hr, followed by washing with DMF. Subsequently, NZ-1 Fab is added onto the grid surface for 45 minutes and then washed by Tris buffer to terminate the crosslink reaction. (C) Procedure for on-grid purification of target proteins form cell lysates by IAAG grid and cryo-freezing. The freshly assembled IAAG grids were immersed in cell lysates containing PA-tagged target for 20 minutes, followed by washing three times with protein buffer to removenonspecifically bound proteins. The grids were immediately cryo-frozen.

### IAAG grid assembly and surface property evaluations

To assemble the IAAG grid, we first modified the GO grid by immersing the GO-coated grid in the Pyr-NHS solution for 1 hour, followed by washing by Dimethylformamide (DMF) (Fig. 1B). We then used X-ray photoelectron spectroscopy (XPS) to assess the efficiency of Pyr-NHS modification on the GO film. The elemental analysis of Nitrogen N 1s from the XPS data revealed a new component emerged on the Pyr-NHS treated GO surface compared with the untreated one (Fig. 2A). This observation indicates successful loading of Pyr-NHS onto the GO film.

**Fig. 2.**
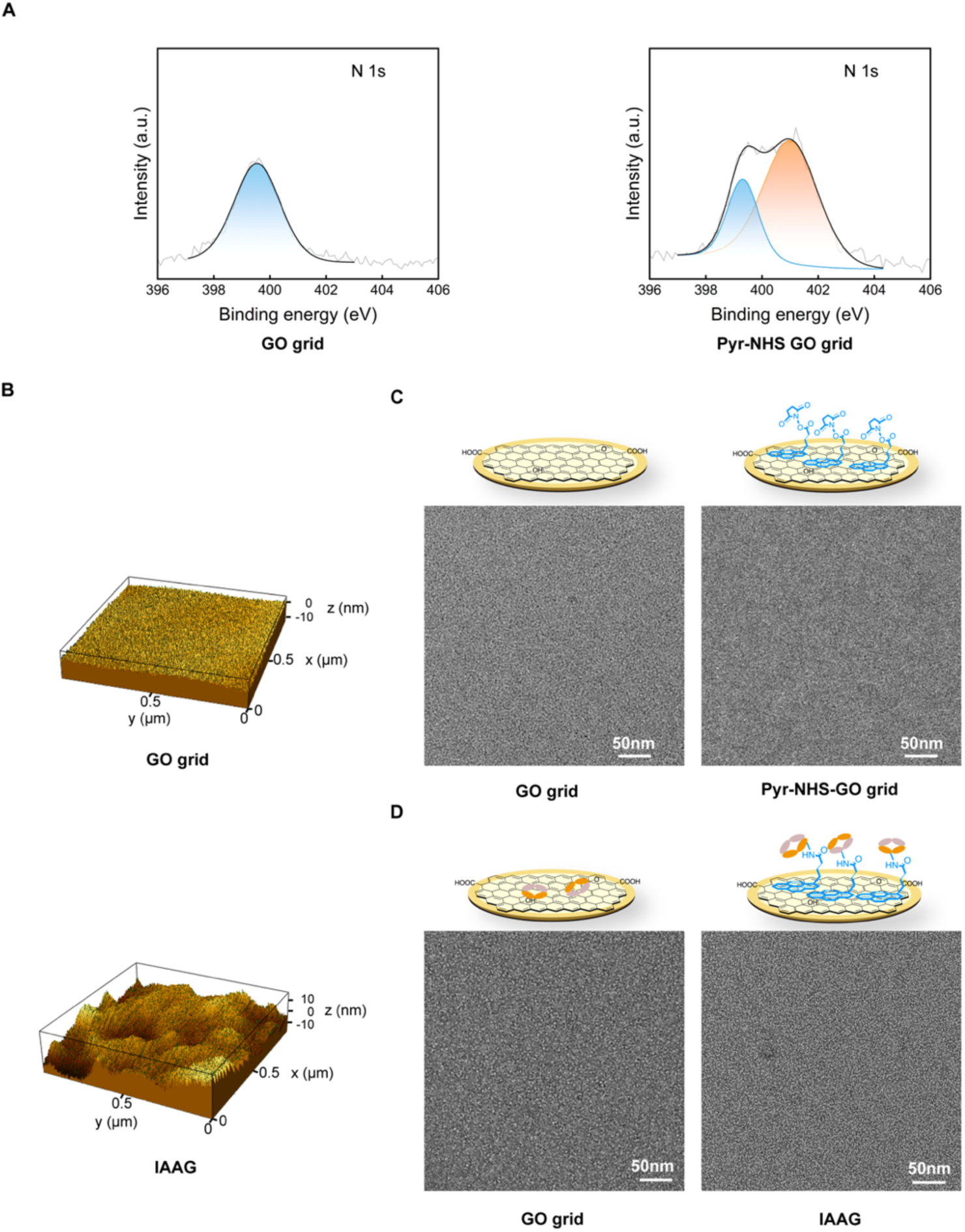
Surface property evaluation of the IAAG grid. (A) XPS spectra of the N 1s of the GO grid (left) and the Pyr-NHS treated GO grid (right). The light gray line represents the original data while the black line presents the fit. The blue peak represents the N 1s state of the untreated commercial GO grid, and the orange color scheme represents the new bond state of N 1s on the Pyr-NHS treated GO surface. (B) AFM mapping of the GO grid (top) and the IAAG grid (bottom). (C) NS-EM image of GO grid (left) and Pyr-NHS treated GO grid (right). (D) NS-EM image of the GO grid (left) and the IAAG grid (right) treated with NZ-1 Fab. It appears that much more Fab densities (small white dots) can be observed on the IAAG grid.

Subsequently, we applied 75 μg/ml NZ-1 Fab onto the Pyr-NHS-treated GO grid and incubated it at room temperature for 45 minutes. The grid was then washed with Tris buffer to terminate the crosslink reaction (Fig. 1B). Atomic force microscopy (AFM) scanning of the grid surface revealed that the IAAG grid surface became apparently more uneven compared with the control GO surface (Fig. 2B), indicating successful immobilization of NZ-1 Fab on the IAAG surface. Additionally, we examined the Pyr-NHS treated GO grid and the control untreated GO grid before and after the application of NZ-1 Fab, using negative staining electron microscopy (NS-EM). This analysis showed that both grid surfaces appeared plain prior to the addition of NZ-1 Fab (Fig. 2C). In contrast, post-application, the Pyr-NHS modified GO grid displayed abundant immobilized NZ-1 Fab (Fig. 2D), while the control grid showed very sparse distribution of Fab. These data collectively indicate the high immobilization efficiency of Pyr-NHS to NZ-1 Fab on the GO surface through crosslinking reaction.

### IAAG can enrich target protein and overcome preferred orientation problems for atomic resolution cryo-EM

Low concentration and preferred orientation problems often impede high-resolution cryo-EM for macromolecular complexes^9,22,52^. To tackle these challenges, we first examined whether the IAAG strategy could enrich low concentration target proteins. We used a low concentration of purified PA-tagged apoferritin, with the PA tag inserted at the exposed N-terminus of apoferritin, as a testing sample. This was evident from the sparse presence of apoferritin on the control GO grid (Fig. 3A). In contrast, we observed an obviously higher quantity of apoferritin on the IAAG grid (Fig. 3B), enabling us to determine an apoferritin cryo-EM structure at 2.5 Å resolution (Fig. 3C, Fig. S1A-C, Table. S1). These data demonstrate that using the IAAG strategy can effectively enrich target proteins for atomic resolution cryo-EM structural determination.

**Fig. 3.**
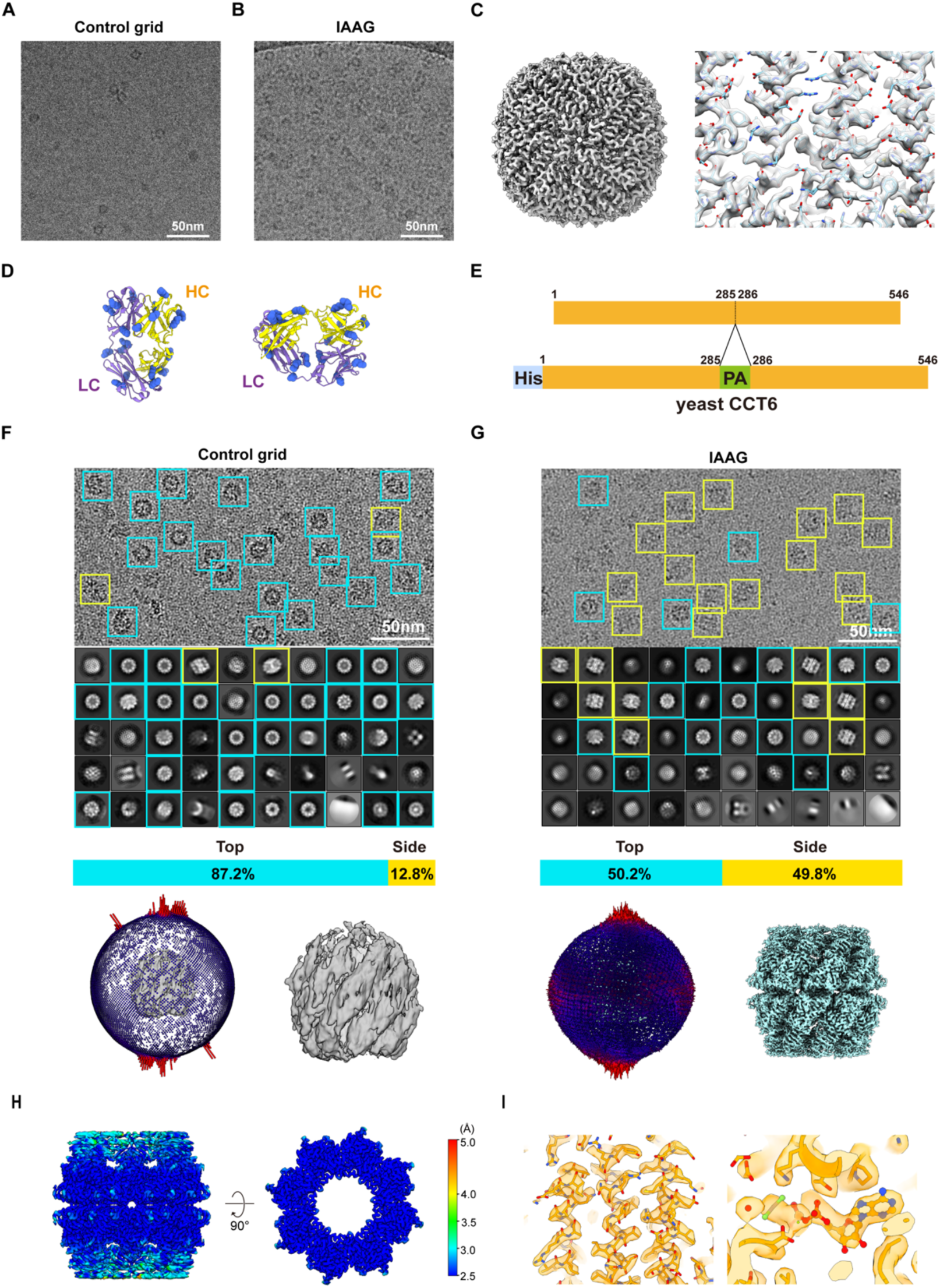
IAAG can enrich target protein and overcome preferred orientation problems for atomic resolution cryo-EM. (A) Cryo-EM image of the PA-tagged apoferritin prepared by control GO grid, showing the low concentration of the target. (B) Cryo-EM image of the same sample prepared with IAAG grid, suggesting enrichment of the target protein. (C) 2.5-Å-resolution cryo-EM map of the IAAG enriched apoferritin (left), and high-resolution structure features (right) (model PDB: 6Z6U). (D) Side and top views of the NZ-1 Fab structure (PDB: 4YNY), with the lysine residues colored in cornflower blue. (E) Diagram of the His tag and PA tag inserted yeast CCT6 plasmid construct. (F) Cryo-EM image, reference-free 2D class averages, top/side view particle population analysis, and the corresponding 3D reconstruction (no imposed symmetry) and particle Euler angle distribution for the CCT6-HR-ATP-AlFx on control Quantifoil grid. This demonstrated obvious preferred top-view orientation problems. Top/tilted top views indicated by cyan square, and side/tilted side views in yellow square. (G) Similar analyses as in (F) for the CCT6-HR-ATP-AlFx on the IAAG grid, which helps overcome the preferred orientation problems. (H) Local resolution estimation of the CCT6-HR-ATP-AlFx (with D8 symmetry), displayed are the side view (left) and central slice top view (right). (I) High resolution structure features (left) and the zoom-in view of the nucleotide pocket (right) (model PDB: 5GW5 subunit CCT6).

Pyr-NHS immobilizes NZ-1 Fab (or other antibodies/Fabs) by cross-linking with the primary amine in the lysine residue or their N-terminus. The presence of multiple lysine residues distributed on the outer surface of NZ-1 Fab (Fig. 3D) allows Pyr-NHS to immobilize NZ-1 Fab in various orientations. Therefore, we sought to test whether our IAAG strategy could help overcome the preferred orientation problems. Our recent study showed that the CCT6 subunit of the yeast group II chaperonin TRiC/CCT can form a homo-oligomeric ring (HR) complex, which exhibits obvious preferred orientation problems for cryo-EM study^53^. A PA tag was inserted into a surface-exposed loop in the middle portion of CCT6 (Fig. 3E). It has been shown that the PA peptide forms a β-turn in the antigen-binding pocket of the NZ-1 antibody^39^. This unique conformation allows the PA peptide to be inserted into various exposed turn-forming loops of target proteins. Our control test on CCT6-HR confirmed the dominant distribution of top view particles (∼87.2%), but a severe deficiency of side views (12.8%) (Fig. 3F). The 3D reconstruction confirmed this preferred orientation problem, which results in a map displaying severe stretch features (Fig. 3F).

We then investigated whether the IAAG strategy could help overcome the preferred orientation problem associated with CCT6-HR. The raw image and reference-free 2D classification result of the IAAG data displayed various views, particularly the previously missing side and tilted side views (which increased to 49.8%) (Fig. 3G). The Euler angle distribution showed the particles distributed more rationally (Fig. 3G), and we determined the cryo-EM structure of CCT6-HR to 2.6 Å resolution (Fig. 3H-I, Fig. S1D-E, Table. S1). These tests collectively demonstrate that our IAAG strategy can effectively overcome preferred orientation problems for atomic resolution cryo-EM. This is likely achieved through the random orientation distribution of the immobilized NZ-1 Fab via the Pyr-NHS linker, and the reduction of AWI interference through the anchoring of the target protein near the GO surface.

### IAAG strategy for on-grid purification of TBCA-apoferritin from cell lysates for high-resolution cryo-EM

Low-abundance and fragile macromolecular complexes often suffer from tedious and time-consuming purification processes for structure studies. The ultimate goal of affinity grid strategy is to provide an efficient and robust method for on-grid purification of target proteins from cell lysates for high-resolution cryo-EM, especially for those rare and fragile biomolecular assemblies. In this study, we aim to examine whether our IAAG strategy can on-grid purify tag-inserted target complexes from cell lysates and assess its feasibility for high-resolution cryo-EM.

Moreover, to address the challenges of small proteins (< 30 kDa) for cryo-EM, strategies have been developed to fuse small proteins to a cage-like structure, such as apoferritin^54^ or DARPin cage^55–57^. We constructed a PA-tagged TBCA-apoferritin display system. This system features a PA tag fused to the N-terminus of TBCA and an apoferritin fused to its C-terminus (Fig. 4A). TBCA, a small protein of 13 kDa, serves as a cofactor that assists tubulin assemble into microtubules. Our SDS-PAGE and NS-EM analyses suggested that the cell lysates of PA-tagged TBCA-apoferritin contain the target complex along with numerous impurity proteins (Fig. S2A-B). To perform on-grid purification (Fig. 1C), we immersed the freshly assembled IAAG grids in the cell lysates containing PA-tagged TBCA-apoferritin for 20 minutes, allowing the targets to bind to the grids. Subsequently, after washing the grids three times with protein buffer to remove nonspecifically bound proteins, we immediately cryo-froze the IAAG grid with on-grid purified samples, which is ready for cryo-EM data acquisition.

**Fig. 4.**
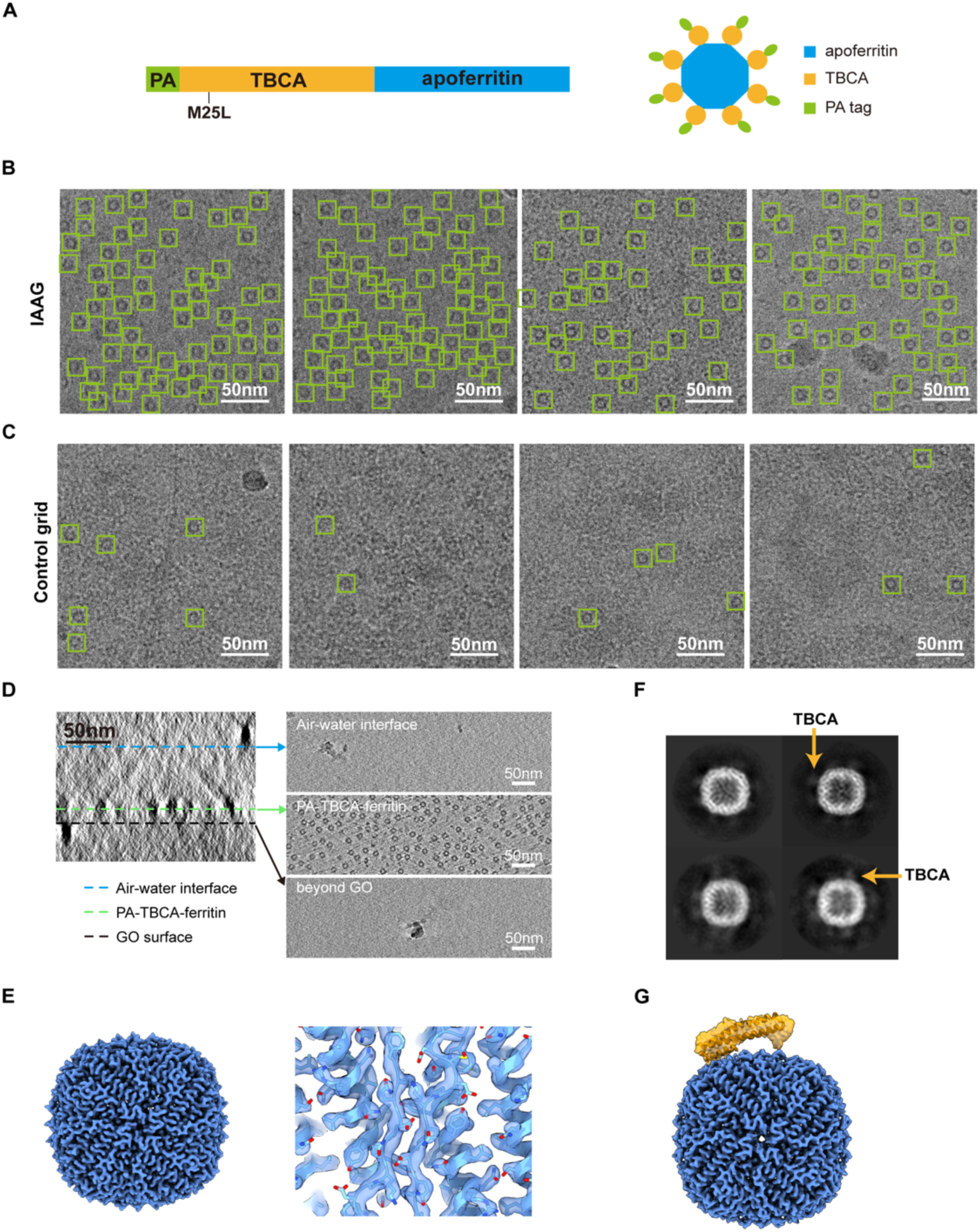
Application of IAAG strategy in on-grid purification of TBCA-apoferritin from cell lysates for atomic-resolution cryo-EM. (A) A diagram of the PA-TBCA-apoferritin plasmid construct. (B) Cryo-EM images of the IAAG on-grid purified PA-TBCA-apoferritin from cell lysates. Many targets (green squire) can be observed. (C) Cryo-EM images of the control GO grid treated with the cell lysates. Only very sparsely distributed target can be observed. (D) Cryo-ET analysis of the IAAG cryo sample, revealing that the TBCA-apoferritin particles are mostly distributed in a layer near the NZ-1 Fab treated GO surface and away from the air-water interface. The AWI is distinguished by the ice contaminations. (E) The 2.4-Å-resolution consensus map of the on-grid purified TBCA-apoferritin, showing the well-resolved apoferritin scaffold (left), and its high-resolution structure features (right, PDB: 6Z6U). For the consensus map rendered in lower threshold also showing the TBCA density, please refer to Fig. S2C. Due to the relative dynamics to apoferritin, TBCA cannot be seen in this higher rendering threshold. (F) 2D class averages of TBCA-apoferritin, which displays fuzzy TBCA density (indicated by orange arrow) around apoferritin. (G) TBCA-focus refined map of TBCA-apoferritin. The TBCA portion (in orange, model from PDB: 1H7C) is rendered at lower threshold (0.04), and apoferritin scaffold (in blue) at higher threshold (0.15).

It appears that the IAAG strategy can effectively purify and enrich PA-tagged TBCA-apoferritin from cell lysates (Fig. 4B). In contrast, the control GO grid failed to do so, displaying only sparsely distributed apoferritin and numerous contaminant proteins from cell lysates (Fig. 4C). To analyze the distribution of TBCA-apoferritin in ice, we performed cryo-electron tomography (cryo-ET) analysis on the IAAG sample. The immobilized NZ-1 Fab anchored target proteins near the GO grid surface (Fig. 4D, Movie S1). The interaction between Fab and target proteins keeps the proteins away from the AWI, thereby avoiding protein denaturation and/or preferred orientation problems in the vitrification process. This demonstrates the potential of the IAAG strategy for overcoming common challenges in cryo-EM sample preparation.

We subsequently determined the cryo-EM structure of TBCA-apoferritin on-grid purified using the IAAG strategy, achieving a nominal resolution of 2.4 Å (Fig. S2C-D, Table. S1). The apoferritin portion was better resolved, displaying atomic resolution structural features (Fig. 4E). This was also confirmed by the local resolution estimation (Fig. S2C), demonstrating that IAAG on-grid purification is feasible for atomic resolution cryo-EM. Still, the exposed TBCA around apoferritin appears very fussy/dynamics in the reference-free 2D class averages (Fig. 4F). As a result, the displayed TBCA was less well resolved (Fig. 4G, Fig. S2E). Further engineering to link TBCA with the apoferritin scaffold could constrain its movements, facilizing its high-resolution structure determination. Putting together, our data suggest that the IAAG strategy is feasible for on-grid purification of target biomolecules from cell lysates and for atomic resolution cryo-EM. When combined with apoferritin display strategy, it has the potential to determine the structure of small proteins such as TBCA.

Pyr-NHS noncovalently anchors onto surfaces sharing sp^2^-hybridized carbon lattice via π-π interaction, which enables the IAAG strategy to be also applied on carbon/graphene surfaces. We tested this by using a continuous carbon-covered grid to assemble the IAAG. Our test demonstrated that the IAAG grid using carbon supporting surface can successfully purify PA-tagged TBCA-apoferritin from cell lysates for cryo-EM (Fig. S3A), while the control grid failed to do so (Fig. S3B). Therefore, carbon/graphene/GO surfaces, all of which share the sp^2^-hybridized carbon lattice, are suitable for assembling IAAG grids.

### IAAG strategy for on-grid purification of CCT6-HR from cell lysates

We further evaluated the on-grid purification ability of IAAG strategy when deal with target complexes with low expression levels. We used yeast CCT6-HR complex as a testing system (Fig. S4A), which expression level in *E.coli* was rather low. The fractions containing the CCT6-HR were barely identifiable in the Coomassie blue-stained SDS-PAGE result (Fig. S4B), and were hardly detected in the NS-EM (Fig. S4C). The ring-shaped CCT6-HR complex was only sparsely observed in fractions 20-22 (Fig. S4C), which were collected for on-grid purification using the IAAG strategy.

Due to the low abundance of CCT6-HR in cell lysates, we extended the incubation time of IAAG grid in cell lysates to 1 hour. In the cryo-EM raw image, a significant number of CCT6-HR in distinct top/side views can be recognized from the IAAG sample, while barely any CCT6-HR could be identified in the control GO grid (Fig. 5A-B). This demonstrates the ability of the IAAG strategy in on-grid purification of low-abundance proteins in cell lysates. We determined the cryo-EM structure of CCT6-HR to a resolution of 2.8 Å using 200 kV electron microscopy data (Fig. 5C-E, Fig. S4D-E, Table. S1). The structure showed high-resolution structure features, with the equatorial domain better resolved (Fig. 5D-E). Also, the Euler angle distribution suggested a reasonable particle angular distribution (Fig. S4D), effectively mitigating the issue of preferred orientation often associated with this system. This underscores the effectiveness of our IAAG strategy in on-grid purification of low-abundance target complexes and addressing common challenges in cryo-EM sample preparation.

**Fig. 5.**
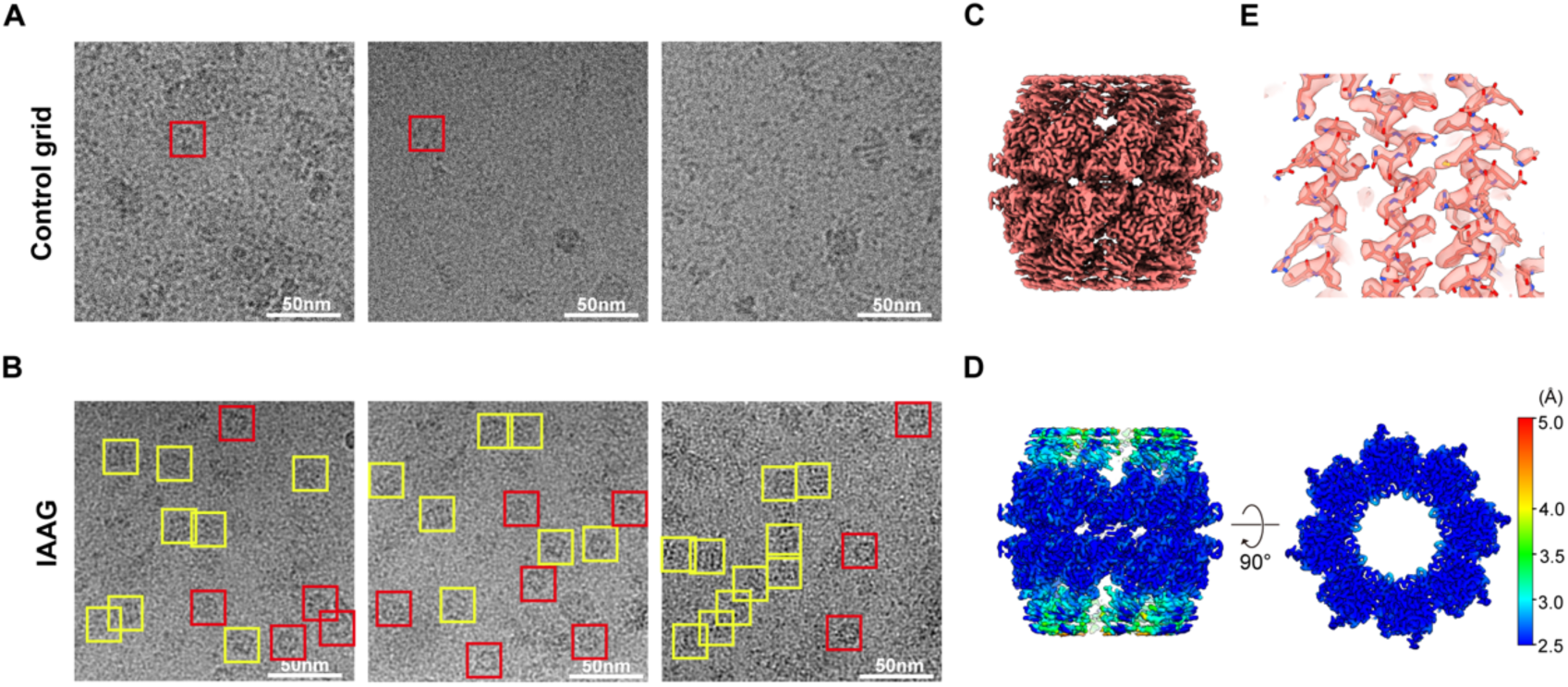
IAAG on-grid purified CCT6-HR from cell lysates for high resolution cryo-EM. (A) Cryo-EM images of the control GO grid treated with CCT6-HR cell lysates, indicating barely any CCT6-HR could be identified in the control grid. (B) Cryo-EM images of the IAAG on-grid purified CCT6-HR from cell lysates. Obviously more number of CCT6-HR in distinct top/side views can be recognized. Here top view indicated by red square, and side view in yellow square. (C) The 2.8-Å-resolution cryo-EM map of CCT6-HR reconstructed from the IAAG data. (D) Local resolution estimation of the CCT6-HR complex. (E) High-resolution structure features of the CCT6-HR complex.

## Discussion

In cryo-EM studies, there are biomolecular assemblies that are extremely low abundance or fragile, and cannot withstand time-consuming and harsh purification or the AWI-related interference in the “standard” cryo-EM sample preparation procedures. For such systems, on-grid purification of target complexes from cell lysates using affinity grids, followed by immediate vitrification, could be powerful solution to for these issues. We developed an efficient immobilized antibody-based affinity grids strategy (Fig. 1). This strategy employs Pyr-NHS as a linker to noncovalently anchor antibody/Fab or bait proteins onto the carbon lattice of the grid surface, including GO, graphene, or carbon, through cross-linking with their lysine residues or N-terminus. We showed that the newly developed IAAG strategy can enrich target proteins (Fig. 3A-B), and overcome preferred orientation problems (Fig. 3F-G). Importantly, it can be used for on-grid purification of low-abundance target complexes (Fig. 1, 5), streamlining the purification process, reducing the demand for large amounts of biological samples, and enabling atomic resolution cryo-EM (Fig. 4, 5). Collectively, our IAAG strategy provides an efficient and robust method for combined sample purification and vitrification, making it feasible for high-resolution cryo-EM.

Contrary to available affinity grid strategies that primarily use freshly homemade graphene or GO supporting film functionalized with customized chemical reagents^15,17,34–37^, our IAAG strategy uses commercially available GO/carbon/graphene grids, and the cross-linker Pyr-NHS is also commercially available. The IAAG grid assembly typically takes less than 2 hours (Fig. 1B). Thus, our strategy is more user-friendly, efficient, and broadly applicable, making it accessible to non-specialist laboratories.

Unlike previous methods that merely place Fab/antibody on the grid surface^16,30,31^, the IAAG strategy immobilizes it through cross-linking with Pyr-NHS (Fig. 1). This enhances stability and effectiveness, allowing IAAG to withstand multiple rounds of grid washing, immersion in cell lysates, and vitrification. Moreover, combined with the high affinity and slow dissociation kinetics of PA tag/NZ-1 antibody^38^, IAAG facilitates on-grid purification of low-abundance target proteins from cell lysates. The presence of multiple lysine residues randomly distributed on the Fab/antibody surface (Fig. 3D) allows it to anchor on the grid surface in various orientations, potentially avoiding preferred orientation problems. Furthermore, the immobilized Fab/antibody keeps the target away from the AWI (Fig. 4D, Movie. S1), avoiding the AWI-related disintegration, aggregation, or preferred orientation issues. It appears that the Fab attached to the grid surface dose not impede atomic resolution cryo-EM structure determinations for apoferritin and CCT6-HR (with the molecular weight of ∼450-1000 kDa). Still, it remains to be tested whether a full-length IgG would generate a background that could impede atomic resolution reconstruction.

Furthermore, the IAAG strategy can utilize other high-affinity pairs, such as an antibody/Fab/nanobody with a specific antigen. More broadly, it has the potential to selectively enrich or on-grid purify target proteins directly from endogenous resources, guided by a specific antibody or other high-affinity binding bait protein. This eliminates the need for special engineering on target proteins. As such, the IAAG strategy could serve as an alternative for cryo sample preparation of fragile assemblies. Additionally, the IAAG on-grid purification could potentially preserve weak interaction partners and provide a more complete conformational landscape under endogenous conditions. This information could be lost with conventional preparation processes, which rely on traditional chromatography purification combined with filtration-based concentration steps^11^. We have also integrated the IAAG strategy with apoferritin display, which could be effective for determining the structure of small proteins (Fig. 4). Overall, our IAAG strategy has broader applicability and potential for various applications in cryo-EM.

In summary, we developed the immobilized antibody-based affinity grid (IAAG) strategy. This approach uses Pyr-NHS as a linker to immobilize antibody/Fab/nanobody on carbon/graphene/GO-coated cryo-EM grids, indicating its wide applicability. Our IAAG strategy can be employed for on-grid purification of tagged complexes from cell lysates, enabling atomic resolution cryo-EM. Furthermore, the Pyr-NHS linker can potentially immobilize other antibody/nanobody or bait proteins, allowing for on-grid affinity purification of endogenous target biomolecules or organelles, without the need for an affinity tag. This development paves the way for a more user-friendly and robust approach with broader applicability in cryo-EM structural studies, and potential extension to cryo-ET *in situ* structure studies.

## Acknowledgements

We are grateful to the staffs of the NCPSS Electron Microscopy facility, Database and Computing facility, and Protein Expression and Purification facility for instrument support and technical assistance. We also thank the Shanghai Nanoport, Thermofisher Scientific for their instrument support on Glacios. This work was supported by grants from Strategic Priority Research Program of CAS (XDB37040103), the NSFC (32130056 and 31872714), National Key R&D Program of China (2017YFA0503503), and Shanghai Pilot Program for Basic Research from CAS (JCYJ-SHFY-2022-008).

## Author contributions

Y.C., Q.Z., X.H., and Z.D. developed the IAAG strategy. Q.Z., X.H., Y.W., Q.H., Q.S., and W.J. constructed the plasmids and purified the proteins. Q.Z. and X.H. prepared the cryo-sample, collected the cryo-EM data (X.S. involved in data acquisition on Glacios), and performed the reconstruction. X.M. performed the cryo-ET. Q.Z. and S.Z. performed the XPS. S.Z. and T.C. performed the AFM. Q.Z., X.H. and Y.C. analyzed the data. Q.Z. and Y.C. wrote the manuscript with the input of X.H. and Y.W.

## Data and materials availability

All data presented in this study are available within figures and in Supplementary Information. Cryo-EM maps have been deposited at the Electron Microscopy Data Bank with accession codes EMDB-38143 for apoferritin, EMDB-38142 for CCT6-HR-ATP-AlFx, EMDB-38145 for TBCA-apoferritin consensus map, EMDB-38146 for TBCA-focus refined TBCA-apoferritin map, and EMDB-38147 for CCT6-HR.

---

**Fig. S1.**
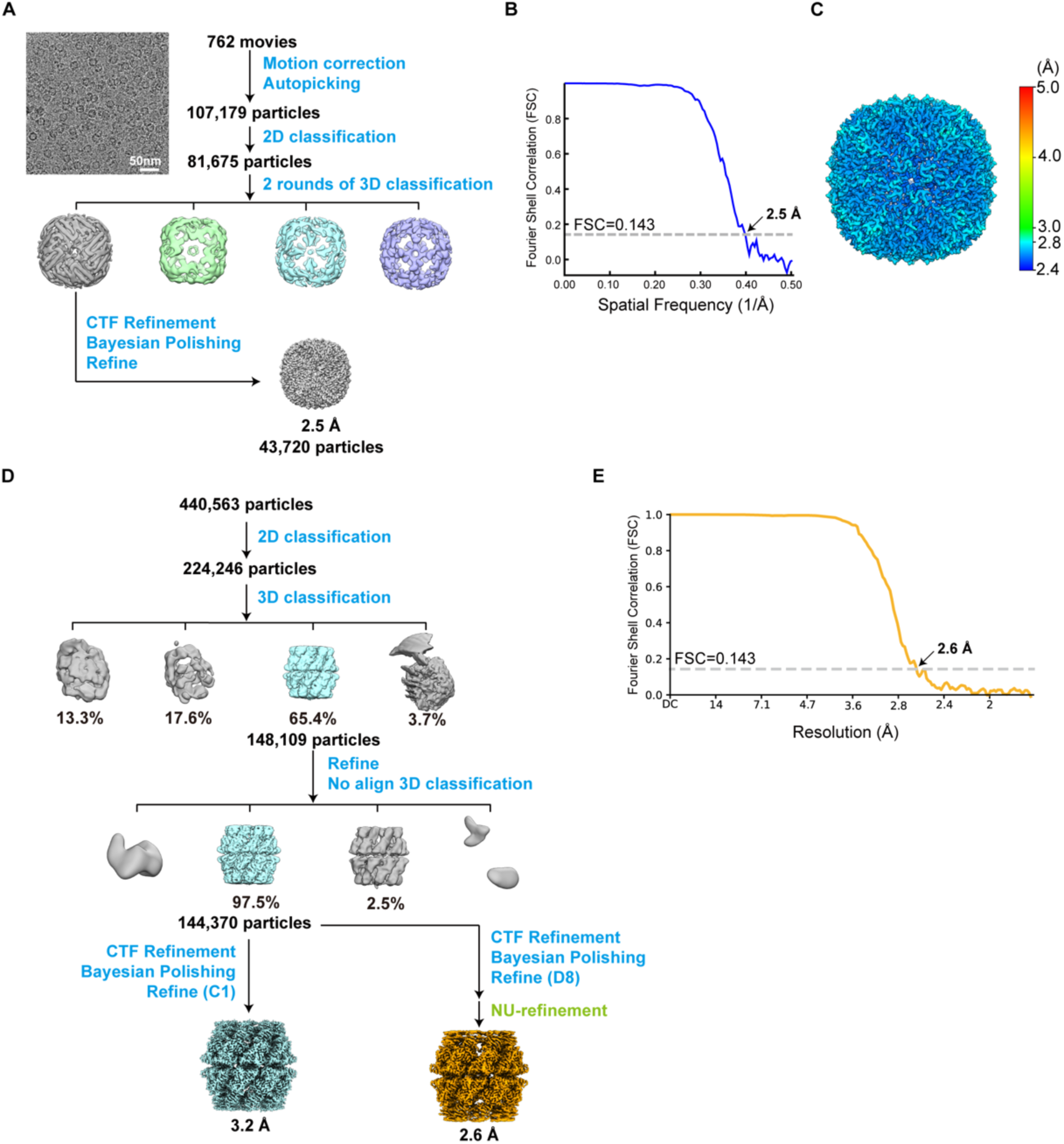
Workflow for the data processing of IAAG-treated apoferritin and CCT6-HR-ATP-AlFx. (A) Workflow for the data processing of IAAG-enriched PA-tagged apoferritin. (B-C) FSC curve (B) and local resolution estimation (C) of the PA-tagged apoferritin. (D) Workflow for the data processing of IAAG treated CCT6-HR-ATP-AlFx. The processes labelled in blue were conducted in RELION 3.1 and those in green in cryoSPARC 4.2.1. This color scheme was applied across all subsequent figures. (E) FSC curve of the CCT6-HR-ATP-AlFx reconstructed with D8 symmetry.

**Fig. S2.**
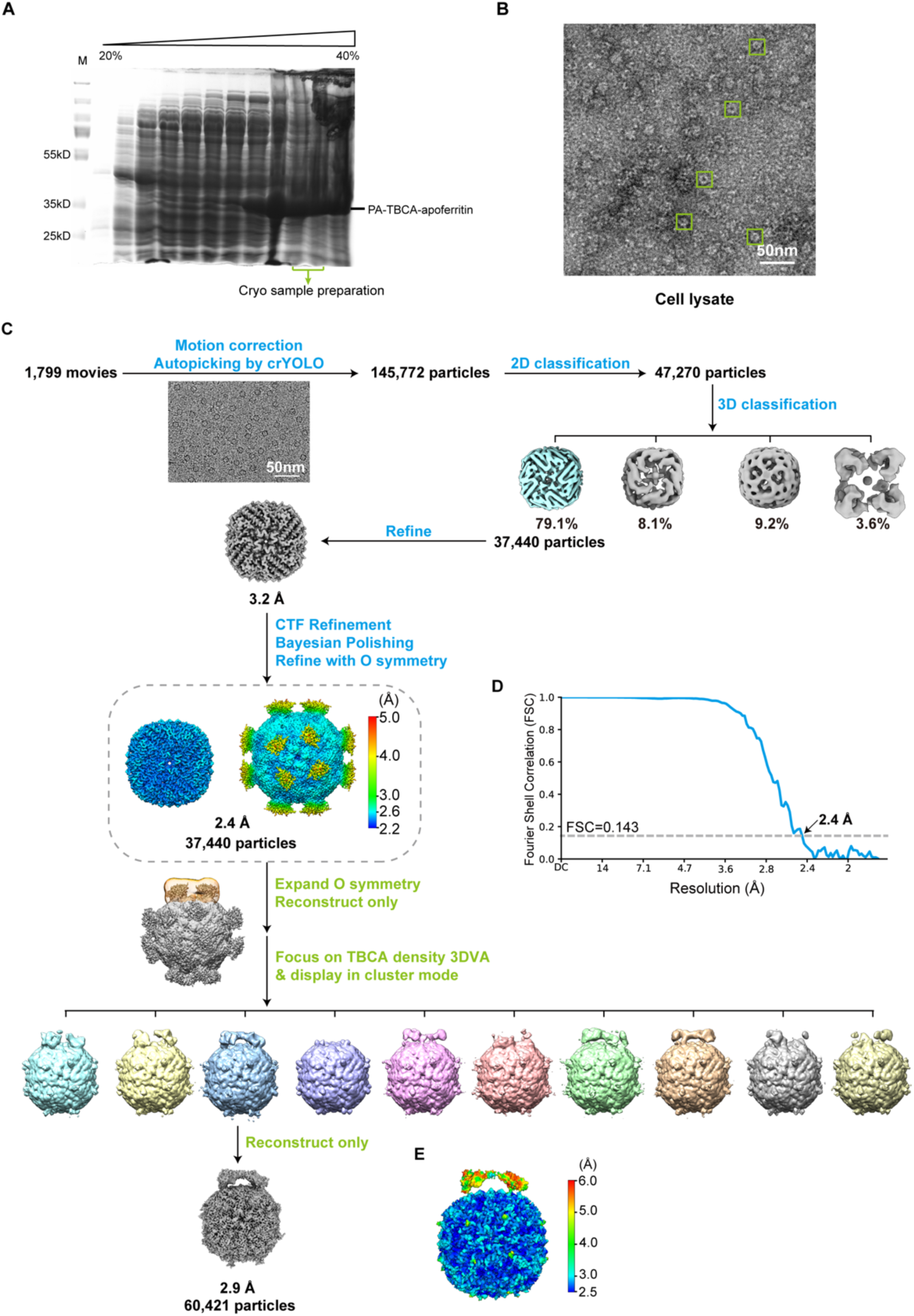
Application of the IAAG strategy in the on-grid purification of TBCA-apoferritin from cell lysates and workflow for cryo-EM data processing. (A) Coomassie blue-stained SDS-PAGE of PA-tagged TBCA-apoferritin cell lysates. (B) NS-EM image of TBCA-apoferritin cell lysates, with the TBCA-apoferritin indicated by green square. (C) Workflow for data processing of the IAAG on-grid purified TBCA-apoferritin. The 2.4-Å-resolution consensus map of TBCA-apoferritin is illustrated within the grey dotted line frame. The local resolution estimation of the map is shown, with thresholds set at 0.15 (left) and 0.015 (right). (D) FSC curve of the consensus map of TBCA-apoferritin. (E) Local resolution estimation of the TBCA-focus refined map of TBCA-apoferritin, displaying improved TBCA density.

**Fig. S3.**
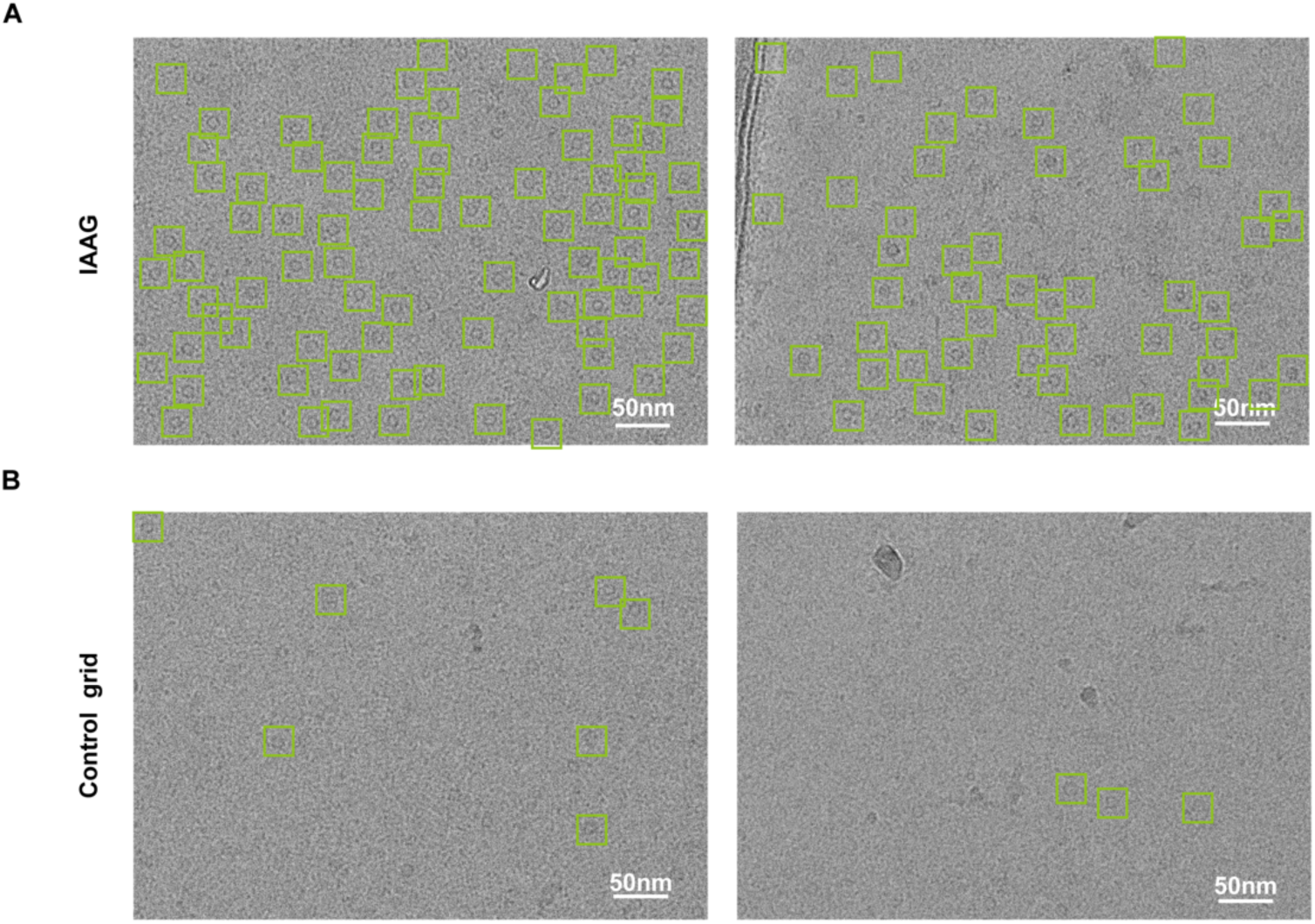
On-grid purification of TBCA-apoferritin using IAAG strategy, assembled with continuous carbon-covered grid. (A) Cryo-EM images of PA-tagged TBCA-apoferritin, on-grid purified from cell lysates using IAAG grid supported by continuous carbon surface. TBCA-apoferritin are indicated with green squares. (B) Cryo-EM images of the control continuous carbon grid treated with cell lysates.

**Fig. S4.**
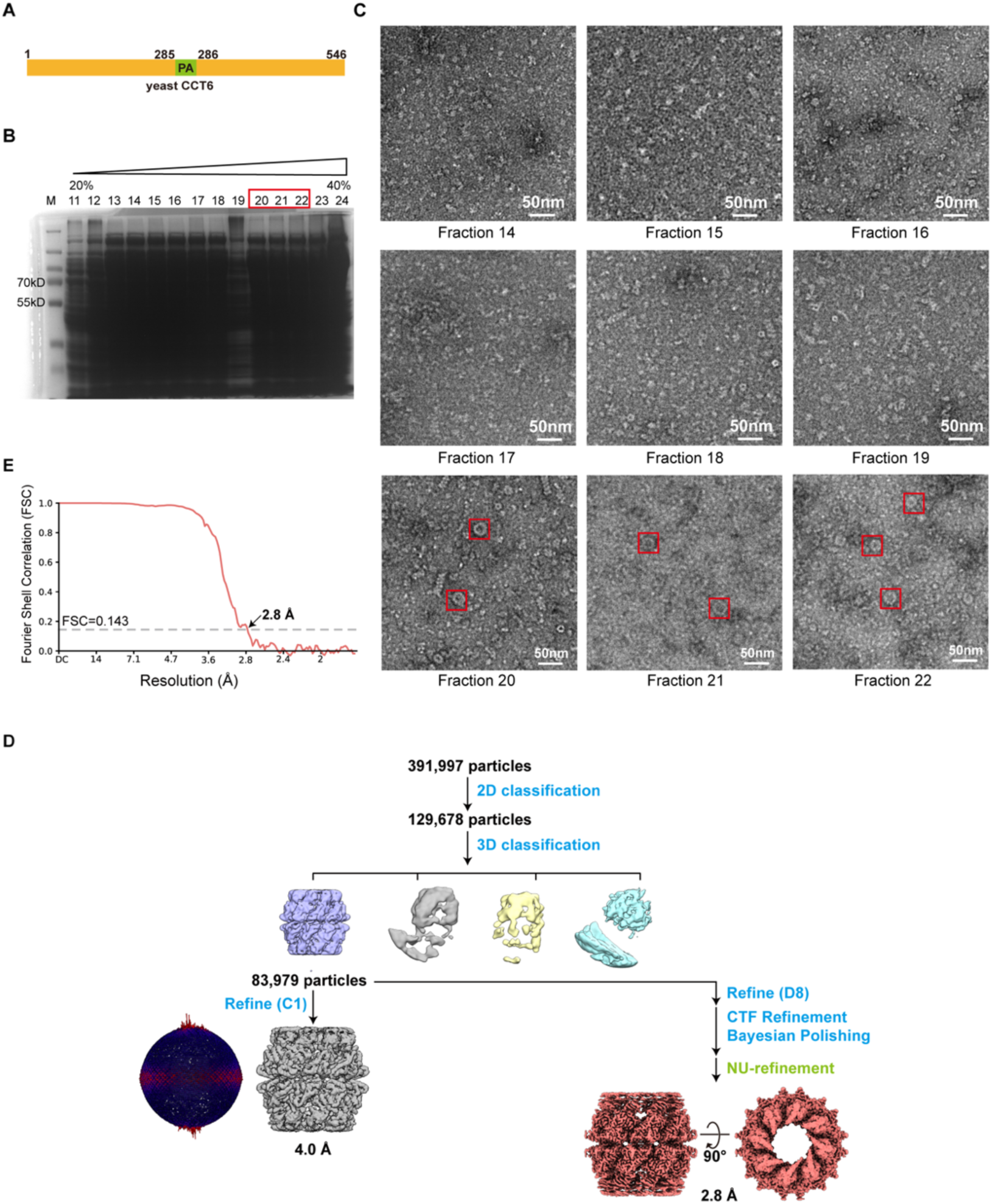
Workflow for data processing of the IAAG on-grid purified CCT6-HR. (A) Diagram of the PA tag inserted yeast CCT6 plasmid construct. (B) Coomassie blue stained SDS-PAGE of CCT6-HR cell lysates. (C) NS-EM images of CCT6-HR crude cell lysate fractions, with most of the fractions showing no observable target. Only in fractions 20-22, very sparsely distributed CCT6-HR (red squire) can be seen. (D) Workflow for the data processing of the IAAG on-grid purified CCT6-HR. Euler angle distribution for the C1 reconstruction is also showed. (E) Resolution assessment by the FSC curve of the CCT6-HR map.

**Table S1.**
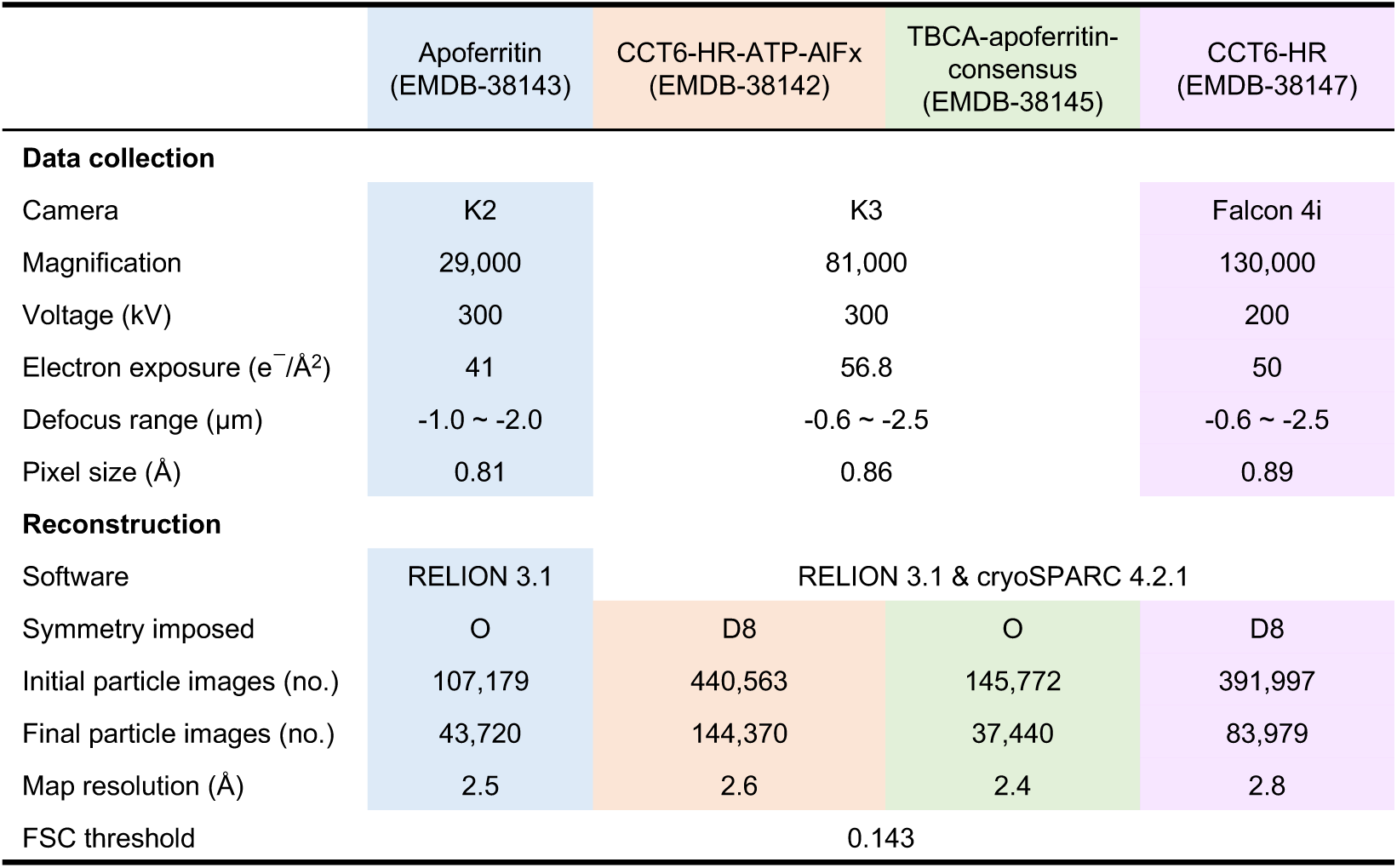
Reconstruction statistics.

**Movie S1. Cryo-electron tomography analysis of the IAAG purified TBCA-apoferritin.** The slices are displayed from the air-water interface to beyond the GO surface. The PA tagged TBCA-apoferritin distributed near the GO grid surface and away from the AWI.

